# Aggregation and Defence

**DOI:** 10.1101/2022.10.25.513673

**Authors:** David M. Durieux, Brad J. Gemmell

**Affiliations:** University of South Florida

## Abstract

*Cassiopea* sp., the Upside-Down Jellyfish, are considered to be sedentary epibenthic organisms which exhibit little to no movement on the seafloor. In this study, we use time-lapse videography of a Cassiopea population in the Florida Keys to demonstrate that Cassiopea sp. exhibit a greater degree of benthic locomotion than previously understood, with animals covering distances up to 178 cm per day. In addition, Cassiopea seem to aggregate on the bottom, with average number of neighbours consistently higher than would be expected for random distributions. Given the ability of Cassiopea to release nematocysts directly into the water column, we present this aggregation as a potential defensive behaviour in this species.

## Introduction

The upside-down jellyfish, genus *Cassiopea* (Péron & Lesueur, 1809) is unique among jellyfish in living an almost exclusively benthic lifestyle, with its bell resting on the bottom and its oral arms extended upwards into the water column. In this position, they pulse their bells in a motion similar to the swimming mechanism of other jellyfish, but rather than displacing their bodies, they create a feeding current which draws prey from the surrounding water into their filter feeding arms. This water is then ejected upwards into a vertical jet with velocities up to 1-2 cm s^− 1^ (Durieux et al., 2021). *Cassiopea sp.* are usually found in dense, patchy aggregations in the wild (Hamner & Dawson, 2009; Keable & Ahyong, 2016). The density of animals can vary greatly, even within a single habitat. For example, in the Red Sea, *Cassiopea sp.* were found to have an average density of 1.6 animals m^−2^, while some areas had up to 31 animals m^−2^ (Niggl & Wild, 2010). Higher densities have been observed in the Florida Keys, up to 97 animals m^−2^ where the median value was 29 animals m^−2^ (Durieux et al., 2021). Nearby regions are often devoid of *Cassiopea sp.*, which suggests a form of aggregating behaviour leading to a patchy distribution. We know that *Cassiopea sp*. are not truly sessile, as they are capable of swimming (Passano, 2004), and changes in their position on the benthos have been observed over periods of days (Jantzen et al., 2010). If *Cassiopea sp*. are able to relocate themselves freely then their dispersion pattern may be of some adaptive significance.

Many species of pelagic jellyfish aggregate in swarms, including the moon jellyfish *Aurelia sp.* (Hamner & Dawson, 2009), as well as the rhizostomeae *Phyllorhiza punctata* (Graham et al., 2003), *Lychnorhiza lucerna* (Brotz et al., 2017), and *Stomolophus meleagris* (Brotz et al., 2017). Some of these aggregations seem to be formed as a result of behavioural patterns of the animals involved, rather than purely physical processes such as water currents. In fact, *Aurelia* have been found to behaviorally compensate for shear flows by altering their bell position and pulse kinematics (Rakow & Graham, 2006). *Aurelia* were also found to swim horizontally towards an existing aggregation, and then alter their behaviour to vertical swimming (Albert, 2011) encouraging the growth of that aggregation, and it is possible that *Cassiopea sp*. do something similar, moving towards conspecifics and settling once they reach the desired location. Alternatively, *Aurelia* have some form of distance sensing, and aggregations of jellyfish maintain a minimum distance from hazardous rock walls and the sea floor (Albert, 2011), which could also concentrate jellyfish to high densities. If *Cassiopea sp.* similarly locate themselves in more favourable habitat, this would naturally lead to aggregations without any drive to be near conspecifics.

Social aggregation is described for other cnidarians such as corals, clonal groups of the anemone *Anthopleura elegantissima* (Ayre & Grosberg, 2005) and the jellyfish *Periphylla periphylla* (Kaartvedt et al., 2015). It is currently unknown whether *Cassiopea sp.* also preferentially aggregate, benefiting in some way from communal living. This could be an effect of fluid dynamics, mating convenience, or defence. On the other hand, aggregation usually leads to competition. We currently lack the information needed to ascertain whether *Cassiopea sp.* preferentially disperse, but are forced close together by limited areas of suitable habitat or by physical mechanisms such as currents. In clonal aggregations the degree of genetic relatedness between individuals may justify this sacrifice, but since *Cassiopea sp.* are not clonal there is likely to be some other trade-off at work, such as defence.

Cnidarians typically defend themselves by means of nematocysts, a diagnostic feature of cnidarians, as they are the only organisms to do so (Fautin, 2009). *Cassiopea andromeda* has been shown to possess five different types of nematocysts: two small isorhizas, two larger birhopaloids, and one type of eurytele (Heins et al., 2015), which are responsible for delivery of venom to both prey and predators. *Cassiopea sp.* exhibit a unique use of these nematocysts for defensive purposes, releasing nematocysts directly into the water column, which has only recently been formally described (Ames et al., 2020). Given that *Cassiopea sp.* move water through their arms, where nematocysts are located and into the water column above in a vertical jet of fluid, this effectively weaponized the water mass above the animal, possibly forming an effective deterrent to predators. Since it would be difficult for a predator to capture a single *Cassiopea sp.* from an aggregation without disturbing neighbouring individuals, it is possible this forms a type of group defence as seen in siphonophores (Mapstone, 2014) and in some anemones (Ayre & Grosberg, 2005).

In this study, we quantify the capacity for benthic locomotion exhibited by *Cassiopea sp.* and explore the potential for *Cassiopea sp.* to organise their aggregations to maximise defensive benefits while minimising competitive costs. We also quantify the amount and time frame of nematocyst release into the water column in the context of aggregations and defence.

## Methods

### Epibenthic Locomotion

During the summers of 2017, 20-21 *Cassiopea sp.* were added to each of three replicate sea-tables at the Keys Marine Laboratory in Layton, Florida, after being collected from the nearby water at this facility. The sea tables (ca. 200 x 80 cm) were filled to a depth of 15 cm seawater that was slowly and continually replaced with new filtered seawater from the nearby bay via an open circuit seawater system. A GoPro HERO2 camera was mounted ca. two metres above each sea table and set to record the entire surface of the sea table by time-lapse photography at one frame per minute for 10 hours. A ruler was placed in each tank for at least three frames for use as a scale. The resulting images, after lens distortion correction, were processed using Fiji image analysis software (Rueden et al., 2017; Schindelin et al., 2012) to manually track the location of each jellyfish. These tracks were then interpreted using a custom Python 3 program. To avoid misinterpretation of tracking errors, any period of sustained displacement greater than 1.5 cm minute^−1^ was considered to be a movement. Average movement duration and distance were calculated, as were average changes in heading. Cumulative distance travelled and displacement from starting position were calculated as well. In addition to the sea table experiments, time-lapse observation of *Cassiopea sp.* in situ was attempted to verify that observed behaviours occurred in the field as well as in captive settings.

### Number of Neighbours

Photographic transects taken in the Winter of 2016, covering 20 quadrats of one m^2^ of benthic surface in a *Cassiopea sp.* aggregation, allowed for visual quantification of *Cassiopea sp.* populations. For each quadrat, the number of each animal’s neighbours, determined as physical contact between the bell or oral arms of two animals, was recorded, and the mean number of neighbours for each quadrat taken, as well as the *Cassiopea sp.* density in that quadrat. For comparison, a simulation was performed using a custom NetLogo (netlogo) program, which simulated random placement of varying densities of *Cassiopea sp.* populations using the known size distribution of a population of *Cassiopea sp.* (durieux2021), and determined the average number of neighbours expected in a randomly-distributed population. The distribution of expected values was then compared to the values observed in situ.

### Nematocyst Release

A five-litre beaker was filled with clean artificial seawater, and a single *Cassiopea sp.* added gently. After allowing the jellyfish to settle for at least 15 minutes, the water was slowly syphoned down to the level of the animal and replaced with clean seawater to remove any nematocysts dislodged during handling. A water sample was taken to record the starting nematocyst concentration. We then agitated the animal for 30 seconds using a glass stir rod, and then collected further water samples at 0.5, 1, 2, 3, 4, and 5 minutes after agitation. The water was then replaced as before, and the animal was allowed to rest for 30 minutes before repeating the treatment. This method was repeated with 6 individual jellyfish. All samples were stained with Lugol’s iodine, and the number of nematocysts in one ml of the sample was counted under a compound microscope at 100X magnification. Mean nematocyst release rates during the first and second agitation events were compared using paired t-tests.

## Results

### Epibenthic Locomotion

Over a 10-hour period, *Cassiopea sp.* were found to engage in significant benthic locomotion (Fig. 1). The maximum recorded velocity over this period was 5.7 mm s-1. The average velocity was 3.8 cm min-1,(± 3.8 SD, n=61), with a maximum of 34.1 cm min-1. Animals spent on average 1.6% of the time moving (± 2.3% SD, n = 61), with a maximum 15.9% of the time. Periods of sustained movement $>$ 1.5 cm were generally short, lasting on average 1.2 min (± 0.6 SD, n = 61) with a maximum observed duration of seven min. Total distance moved was substantial, with average gross displacement of 105.3 cm day-1 (± 64.2 SD, n = 61), with a maximum of 513.5 cm day-1. In terms of net change in position, animals moved on average 46.5 cm day-1 (± 23.6 SD, n = 61), with a maximum of 178.2 cm day-1. Meandering was substantial as well, as absolute values of changes in heading averaged 43.7° (± 42.5° SD, n = 61), with a probability of changing turn direction, from clockwise to anticlockwise or reverse, of 58%. Average turn angle was −0.6° (± 64.0 ° SD), indicating no preference for one direction or another. Observation of populations of *Cassiopea sp.* in the field confirmed that benthic locomotion occurs in situ as well.

**Figure 1.**
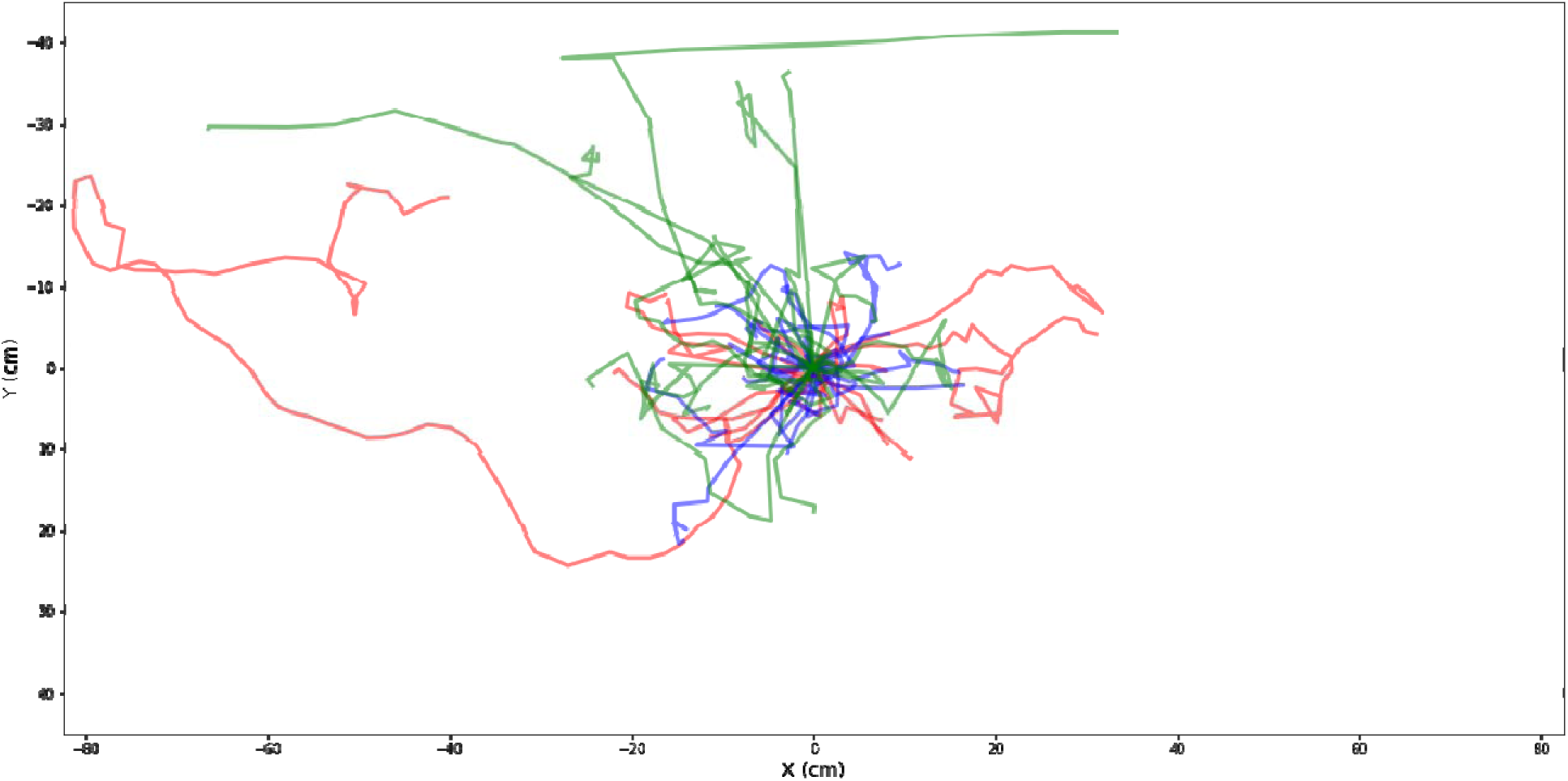
Tracks of 61 *Cassiopea sp*. engaging in benthic locomotion, centred at their starting positions, over a period of 10 hours. Each line represents a single animal, and colour indicates the replicate sea-table in which *Cassiopea sp.* were imaged. The maximum velocity observed was 5.7 mm s^−1^, and the longest track was 2.1 m in length.

### Number of neighbours

Compared to a simulation of randomly-dispersed *Cassiopea sp.* at a variety of population densities (Fig. 2), all but one of the observed quadrats had greater than the mean expected number of neighbours, and 60% of observed quadrats exceeded the 95% confidence interval for the expected number of neighbours. This aligns with in-situ observations of chain-like clusters of *Cassiopea sp.* forming in areas with intermediate population densities (Fig. 2).

**Figure 2.**
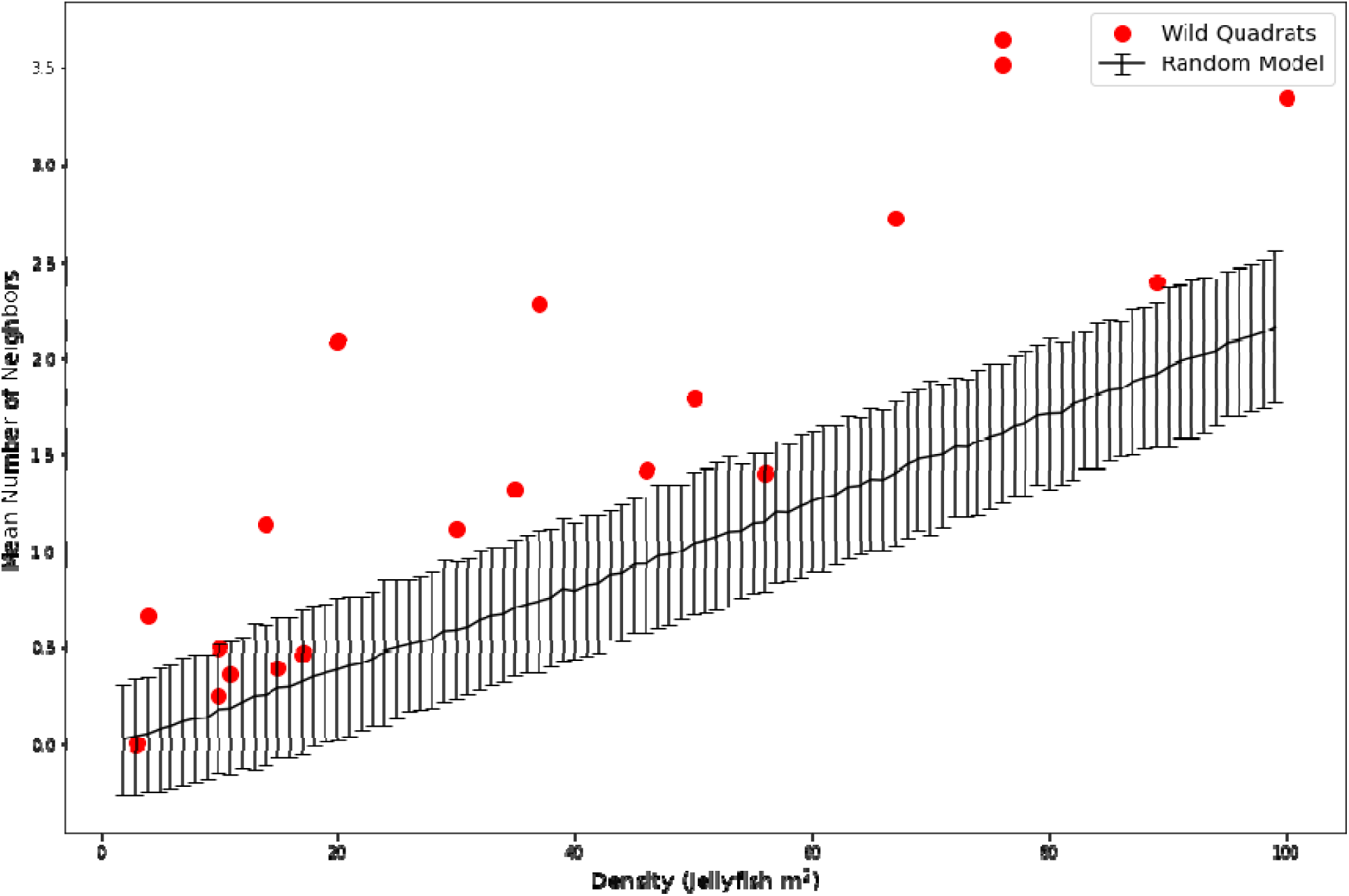
Average number of neighbours plotted against population density for wild *Cassiopea sp.* (Red), and for a simulated randomly dispersed population (Black), where error bars represent the 95% confidence interval of the model. Wild *Cassiopea sp.* have more neighbours relative to population density than predicted by the model in all but one case, and are above the confidence interval in most cases.

**Figure 3.**
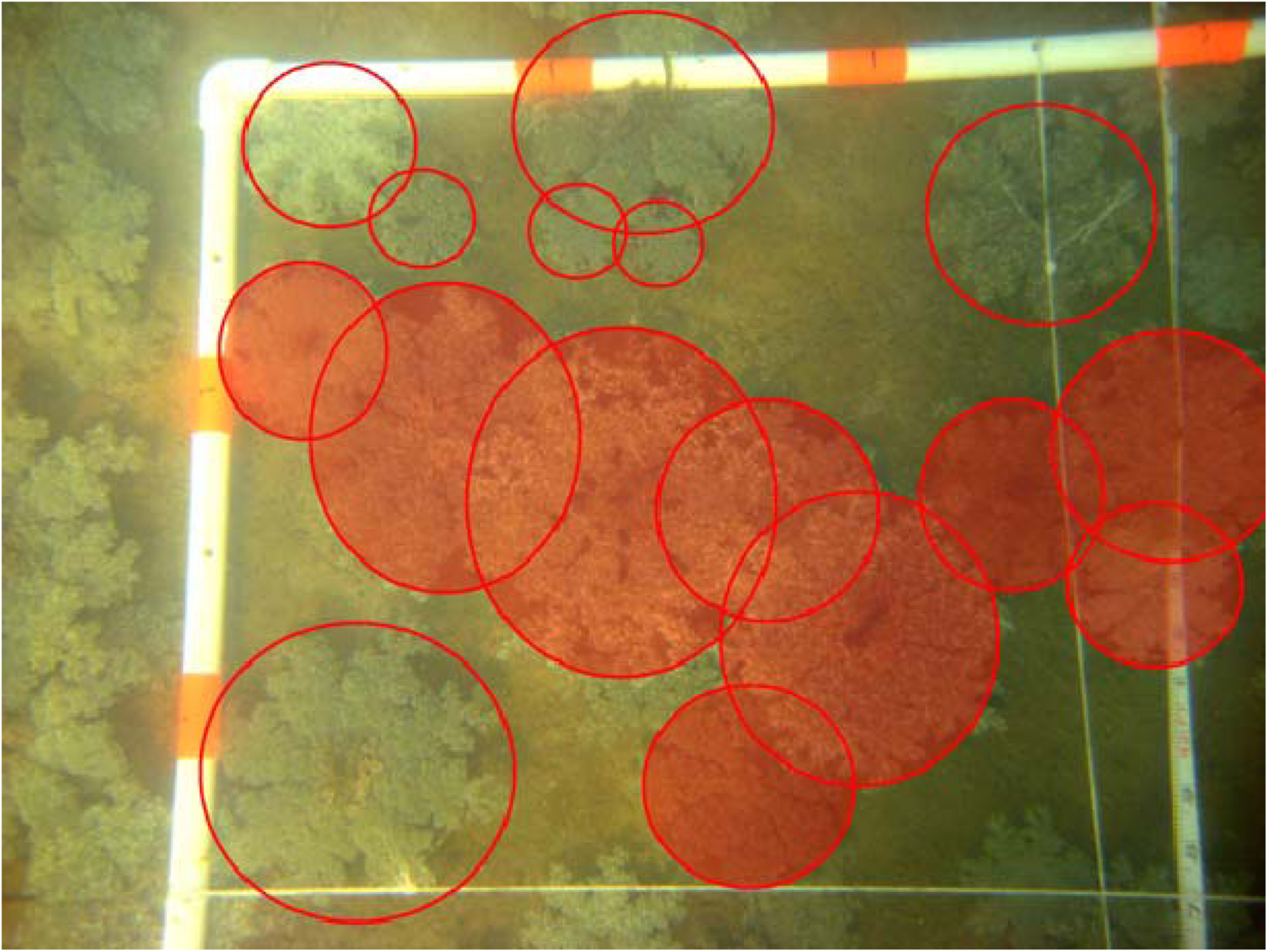
Formations of chain-like clusters of *Cassiopea sp.* were frequently observed in-situ in Layton, Fl.

### Nematocyst Release

Following physical disturbances, *Cassiopea sp.* release significant numbers of nematocysts into the water column (Fig. 4). The initial disturbance event triggered the release rate of on average 43,349 cells cm^−2^ (± 26,510 cells cm^−2^ SD), while a repeated disturbance 30 minutes later caused a mean release rate of 34,755 cells^−2^ (± 30,461 cells cm^−2^ SD). These nematocyst releases were accompanied by other visually observed behavioural changes in *Cassiopea sp.*, such as an increase in mucus production, increased bell pulsation rate, and retracted oral arms, which were not quantified in this study. Release rates were quantified over time (Fig. 5). Following disturbance, 88% of nematocysts were released during the first 30 seconds following the first disturbance. The second disturbance triggered similar timing, with 81% of cells released during the first 30 seconds. Cells were typically not released more than 60 seconds after disturbance.

**Figure 4.**
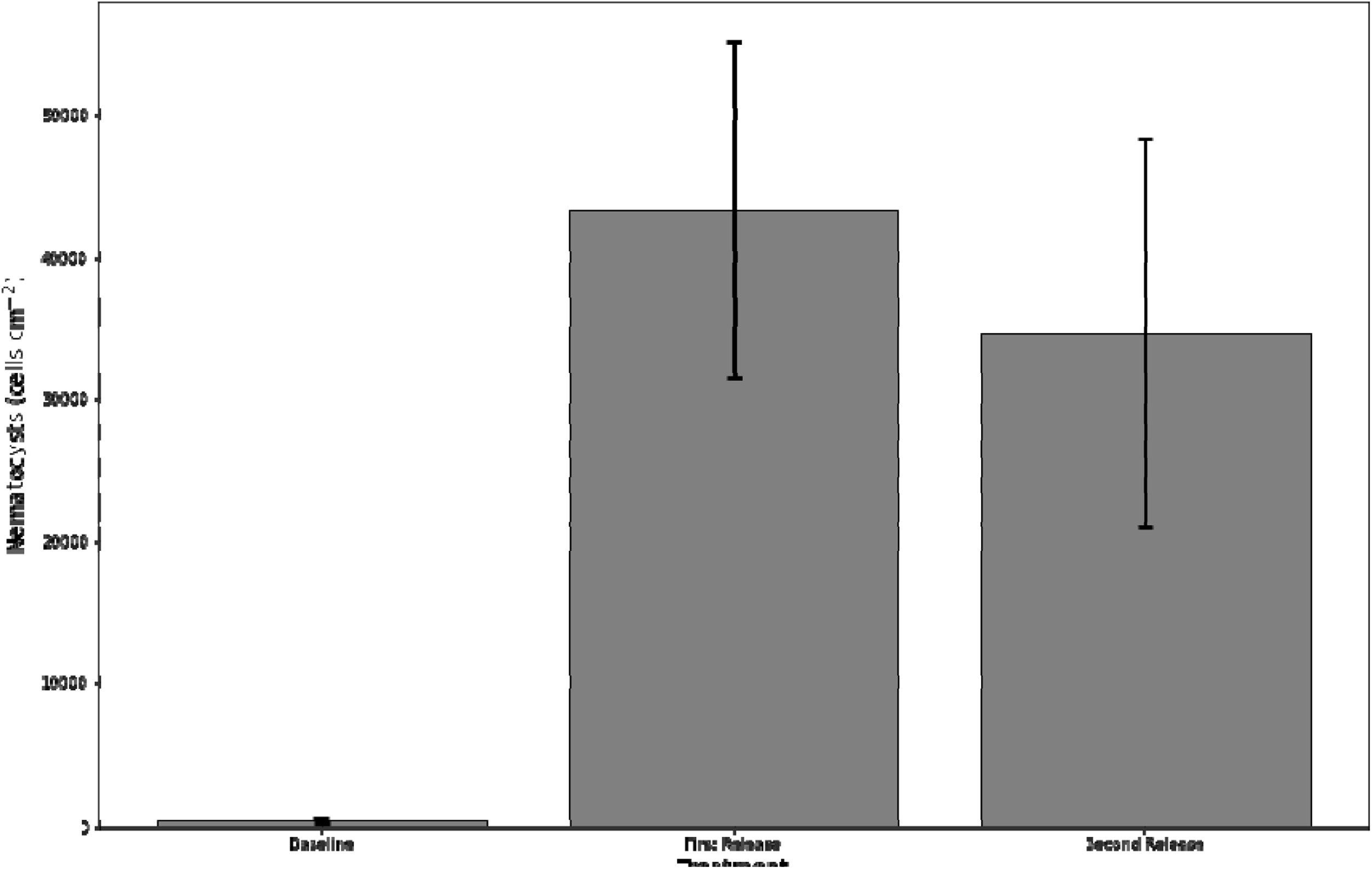
Nematocyst release during the first minute following physical disturbance of *Cassiopea sp.* Mean (± SD) nematocyst counts per cm^2^ of jellyfish bell. Both the first and second agitation events produce releases that are significantly different from baseline (Paired t-test,n=4, p=0.03 and p=0.07 for first and second events respectively), but the two events did not differ significantly from one another (paired t-test,n=5, p=0.33).

**Figure 5.**
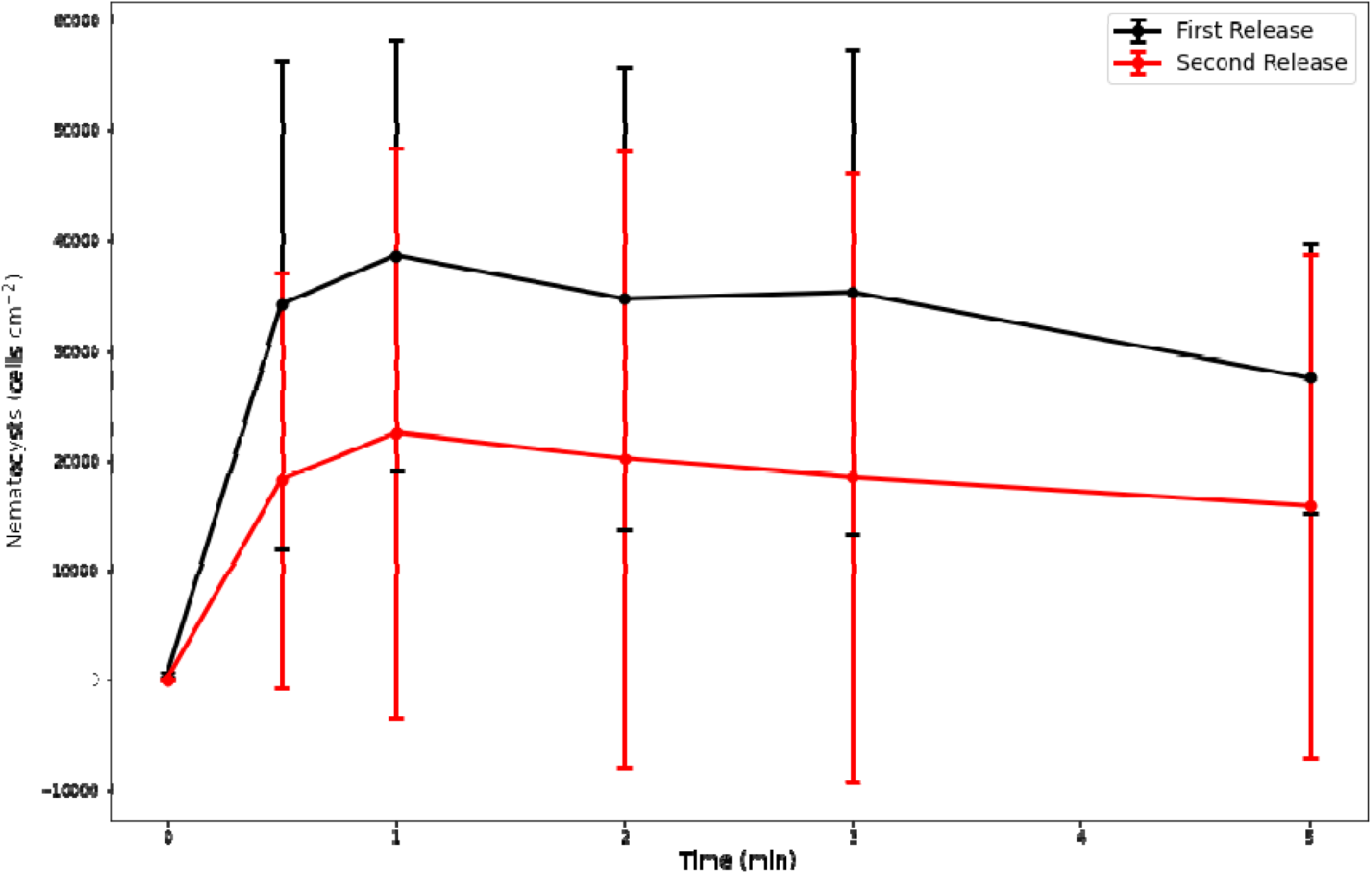
Nematocyst release over time following physical disturbance of *Cassiopea sp.* Mean (± SD) nematocyst counts per cm$^2$ of jellyfish bell after the first (black) and subsequent (red) disturbance events.

## Discussion

One drawback to life as a sedentary bottom-dweller is the reduced ability to respond effectively if one’s habitat becomes inhospitable. Sedentary benthic macrofauna are disproportionately harmed by trawling (de Juan et al., 2007) or storm-induced sediment burial (Abesamis, 2018). While motile organisms are able to simply relocate to a more suitable location, sessile organisms are required to weather difficult conditions. Another condition likely to affect sessile organisms is the formation of hypoxic zones (Vaquer-Sunyer & Duarte, 2011), but it is interesting to note that some jellyfish polyps might actually benefit from hypoxic conditions, as their low oxygen demand allows them to outlast other sessile competitors (Miller & Graham, 2012). While *Cassiopea sp.* polyps typically attach to decomposing mangrove leaves (Fleck & Fitt, 1999) which can be moved by environmental forces, they are not known to actively relocate themselves so these pressures may apply to *Cassiopea sp.* as well.

While adult *Cassiopea sp.* are known to be able to swim, they are generally considered sessile organisms (Gaddam, 2016) and are thus considered to be similar to anemones, which are also capable of relocating themselves if necessary (Muller-Parker & Davy, 2001). However, our results show that *Cassiopea sp.* actually exhibit frequent and active movement across the benthos (Fig. 1). Of 61 animals tracked in this study, on average, *Cassiopea sp.* moved over one m day-1, and spent on average 1.6% of the time actively moving at mean velocities of 3.8 cm min-1. Some individuals moved much more than this, with one animal travelling over five m day-1, indicating a great deal of behavioural variation between individuals. No preference was observed for any particular direction in this experiment. While this slow movement may aid in finding a more favourable position within a mangrove habitat *Cassiopea sp.* require nutrients and must consume benthic prey and/or release interstitial, nutrient-rich porewater (Durieux et al., 2021). Since prey and nutrients could be easily depleted if permanently fixed in one location, one potential use of slow movement across the benthos would be as a means to bolster nutrient or food intake by moving the animal onto fresh habitat that has not been depleted of these resources.

*Cassiopea sp.* are known to occur in clumped distributions (Durieux et al., 2021; Niggl & Wild, 2010), with small individuals aggregating more densely (Tsingalia, 2014). Although this aggregation behaviour is well documented, the mechanism has not been investigated. Benthic locomotion could explain the aggregative mechanism of *Cassiopea sp.* In addition to aggregating into patches, *Cassiopea sp.* also appear to organise themselves spatially within these patches. In particular, they appear to maintain physical contact with neighbours, as the average number of neighbours was significantly higher than would be expected of a random distribution of *Cassiopea sp.* within an aggregation (Fig. 2). This physical contact has been shown to reduce the volumetric flux of the feeding current in *Cassiopea sp.* (Durieux et al., 2021) due to interference between the horizontal incurrent flows of multiple animals, which was aggravated as additional neighbours were added. This is due to the horizontal incurrent flow, which flows along the benthos radially towards the jellyfish (Durieux et al., 2021). The presence of another jellyfish can physically impede this flow. In addition, since this feeding current captures prey from the benthic area surrounding the jellyfish, a reduction of benthic area available to an individual, as well as competing currents from neighbouring animals, can reduce feeding opportunities. This places a negative feedback on aggregation density and makes it likely that some benefit is gained that offsets this competitive effect.

Of course, there are some known benefits to aggregation. For example, aggregating has been shown to improve mate-finding in some species (Ritz et al., 2011; Susset et al., 2018). Another common function of aggregation is predator dilution, where an increase in prey abundance reduces the likelihood of any given prey animal being captured (Aukema & Raffa, 2004). Group defence is another purpose for aggregation, and is known to occur in other cnidarians, such as siphonophores (Mapstone, 2014) and anemones (Ayre & Grosberg, 2005), but both of these cases involve clonal colonies. In spiny lobsters, group defence has been demonstrated to be more common in robust species, which are more capable of defensive behaviour, than in more delicate species (Briones-Fourzán & Lozano-Álvarez, 2008). An individual *Cassiopea sp.* is clearly capable of defensive behaviour. When disturbed, they have been shown to release cassiosomes, complex structures consisting mostly of stinging nematocysts (Ames et al., 2020). We have also observed the release of individual nematocysts, as well as clusters of 2-4 cells. In this study, we demonstrated that *Cassiopea sp.* is capable of releasing in excess of 40,000 stinging nematocysts cm^−2^ into the water column in response to agitation, and are capable of multiple releases (Fig. 4). Such behaviour is unknown outside of *Cassiopea sp.* and a few related rhizostomeae (Ames et al., 2020). This nematocyst release is rapid in response to physical agitation (Fig. 5) and effectively weaponizes the surrounding water column as nematocysts are transported into the water column by the feeding current. The nematocysts themselves can occur clustered in motile units that swim by ciliary action (Ames et al., 2020), further dispersing them around *Cassiopea sp.* It has been observed that *Cassiopea sp.* presence reduced parrotfish grazing on seagrass by 94% (Stoner et al., 2014), indicating that proximity to *Cassiopea sp.* provides a defensive benefit. The same study observed a yellowfin mojarra dying after swimming through *Cassiopea sp.* mucus (Stoner et al., 2014). Since it would be difficult for a predator to capture a single *Cassiopea sp.* within an aggregation without disturbing others in contact with it, a relatively large number of animals could release nematocysts in response to a single predation event.

We have suggested possible benefits and drawbacks to *Cassiopea sp.* aggregation in terms of feeding current inhibition and defence. This creates a trade-off situation where the animal might attempt to balance aggregation and dispersion, and there is evidence to suggest this is the case. Rather than forming tight clusters, *Cassiopea sp.* appear to form chains with one or two neighbours (Fig. 2) at the intermediate densities which allow this. Such an arrangement minimises the reduction of the feeding current while still allowing for a group defence. It appears that *Cassiopea sp.* use their benthic motility to aggregate, and they seem to do so in a manner that allows for group defence benefits when multiple neighbouring animals release nematocysts in response to a disturbance.

